# Accelerating inference in genomic and proteomic foundation models via speculative decoding

**DOI:** 10.64898/2026.01.13.699044

**Authors:** Kimonas Provatas, Aris Karatzikos, Charalampos Koilakos, Michail Patsakis, Alexandros Tzanakakis, Akshatha Nayak, Ioannis Mouratidis, Evangelos Ioannis Avgoulas, Ilias Georgakopoulos-Soares

## Abstract

Genomic and protein foundation models (GFMs and PFMs) have demonstrated strong performance in learning the language of DNA and proteins, but their use in large-scale sequence generation is limited by the latency of autoregressive decoding. Because every token triggers a forward pass of a large Transformer, whose inference is relatively slow, long-sequence generation quickly becomes costly. In this work we adapt speculative decoding to a representative GFM: the DNA model DNAGPT and two representative PFMs: ProGen2 and ProtGPT2. We implement a probabilistic variant of speculative decoding, in which a lightweight draft model proposes short token spans and a larger target model verifies or corrects them in parallel, while preserving the target model’s sampling distribution. Across all three models we systematically study the effect of speculation window length, temperature, draft architecture and prompt length, and we benchmark tokens per second over multiple runs per configuration. Speculative decoding yields consistent speedups over standard key-value cached decoding, with maximum observed speedup reaching 100% increase, while average gains across models ranging between 20% and 40% (e.g. 1.2x–1.4×), without changing the underlying target model predictions. Our results show that speculative decoding is a practical and model-agnostic strategy for accelerating genomic and proteomic sequence generation without sacrificing prediction quality.

## Introduction

Large language models (LLMs) have transformed natural language processing by introducing autoregressive frameworks capable of predicting complex, context-dependent sequences with high accuracy. Genomic LMs (gLMs) extend these autoregressive principles to DNA, enabling the generation of biologically realistic synthetic sequences that capture long-range dependencies, regulatory structure, and evolutionary constraints (Benegas et al. 2025). Protein LMs (pLMs) apply these architectures to learn protein structure and function and to enable tasks such as structure prediction, mutational effect estimation, and *de novo* protein design from sequence (Xiao et al. 2025). As a result, gLMs and pLMs are increasingly used in synthetic biology, functional genomics, and design-driven applications that require accurate modeling of sequence-function relationships (Consens et al. 2025).

Speculative decoding, originally developed to reduce inference latency in AI systems, combines a lightweight draft model that generates multiple candidate tokens with a larger, more accurate target model that verifies them in parallel (Leviathan et al. 2022). This approach substantially accelerates token generation while maintaining model fidelity, enabling near-real-time inference without sacrificing performance. The same principles of predictive parallelism and multi-stage verification are highly relevant for computational biology, where generation of several candidate genomes or proteins is often required to achieve optimal results (Merchant et al. 2025; Madani et al. 2023).

Standard genomic generation using transformer-based language models follows an autoregressive process. Given a sequence of DNA tokens *x*_1+1_, the model predicts the probability distribution for the next token *x*_1+1_. This process is memory-bandwidth bound rather than compute-bound. For every single token generated, the model parameters must be moved from High Bandwidth Memory to the GPU compute units. For a target model *M*_1_ (e.g., DNAGPT-3B (Zhang et al. 2023)), generating a sequence of length *N* requires *N* serial forward passes, resulting in significant latency for long genomic sequences. Similarly, in ProGen2-base (Nijkamp et al. 2023) or ProtGPT2 (Ferruz et al. 2022) architectures, generating a sequence of length *T* requires *T* serial decode steps. Each step is typically memory-bandwidth bound rather than compute-bound, which results in substantial latency for long genomic or proteomic sequences and limits throughput in applications that require large numbers of generations.

Speculative Decoding (Chen et al. 2023; Leviathan et al. 2022) addresses this bottleneck by introducing a smaller, faster draft model *M_d_* that approximates the next-token distribution of the target model *M_t_*. Instead of asking *M_t_* to generate one token at a time, speculative decoding proceeds in two stages. First, the draft model generates a short sequence of *L* proposed tokens, starting from the current prefix. Second, the target model evaluates this entire proposed span in parallel by computing logits for all positions in a single forward pass. The algorithm accepts or rejects each proposed token in a way that guarantees that the final sequence has the same distribution as direct sampling from *M_t_*.

Our work represents the first application of speculative decoding to biological language models. In the probabilistic variant used here, each draft token is accepted with probability equal to the minimum of one and the ratio of its probability under the target and draft models. If a token is rejected, a replacement is sampled from a residual distribution based on the difference between target and draft probabilities. If all proposed tokens in a group are accepted, the algorithm may additionally draw a bonus token directly from the target. This design allows one to amortize a single, expensive forward pass of the target model over multiple token positions, while preserving the statistical behavior of the original model. In our experiments, the maximum observed speedup reached a 100% increase, while average gains across models ranged between 20% and 40% (i.e., 1.2x–1.4x) compared to standard KV-cached autoregressive decoding throughput.

## Methods

### Speculative decoding for gLMs

As a proof of concept, our DNA experiments were based on DNAGPT (Zhang et al. 2023), a decoder-only transformer architecture designed specifically for genomic sequence modeling. DNAGPT is pre-trained from scratch on large-scale genomic datasets, allowing it to capture long-range biological dependencies and complex regulatory motifs. For speculative decoding we employed a dual-model setup consisting of a target model *M_t_* and a draft model *M_d_*. The target model was dna_gpt3b_m, a 3-billion-parameter variant that served as the verification oracle and provided high-fidelity representations of sequence probability. We used dna_gpt0.1b_m, a lightweight variant with 0.1 billion parameters that has been optimized for low-latency inference, as a draft model. Both models shared the same tokenizer and vocabulary, which ensured that they are compatible for speculative decoding.

### Speculative decoding for pLMs

For protein sequence modeling we consider ProGen2 (Nijkamp et al. 2023), a family of protein language models trained on large collections of natural and synthetic proteins, and ProtGPT2 (Ferruz et al. 2022), which has been trained on UniRef data using byte-level BPE over amino-acid characters. For ProGen2, we used the HuggingFace implementations hugohrban/progen2-xlarge as the target model and hugohrban/progen2-small as a pretrained draft model. ProGen2 uses its own tokenizer, which encodes amino-acid sequences along with optional metadata tokens such as tags and separators. In our benchmarks we followed the recommended prompting scheme, including the appropriate tag tokens and using start tokens such as 1M for generation. Sequences were decoded until a ProGen-specific termination token was produced or a fixed maximum number of tokens was reached. Unlike ProGen2, ProtGPT2 has no smaller official sibling model available that could be used as a draft. To apply speculative decoding to ProtGPT2, we therefore constructed all drafts by truncating the Transformer stack of the full model, as described in the next subsection.

### Truncated Draft Models

For ProGen2 and ProtGPT2 we additionally explored truncated draft models that are directly derived from the target. The idea was to reuse the tokenizer, embedding matrix and output head of the target model, while keeping only a subset of the Transformer blocks. Concretely, we located the GPT-2-like stack, which is typically exposed as a module with a list of blocks, …,. We then defined a new Causal LM that retained the token embeddings, the dropout and layer normalization modules, and the final language modeling head, but replaced the full stack of blocks by the first *k* blocks only. We systematically varied the effective depth *k* when testing speculative decoding, especially in the ProtGPT2 experiments where truncated models are the only available drafts. We experimented with using the last layers, intermediate layers, or combinations of initial/intermediate and last layers but the use of initial contiguous layers yielded the best acceptance ratio. The resulting model had the same input and output spaces as the target but a shallower depth, and it supported the same KV-cache interface. These truncated drafts can be viewed as structurally distilled approximations to the target model. They are cheaper per token, because attention and feed-forward operations are executed in fewer layers, yet they preserve much of the target’s representational structure.

### Speculative Decoding Algorithm - Probabilistic Acceptance

Our implementation followed the probabilistic speculative decoding scheme outlined by (Chen et al. 2023; Leviathan et al. 2022). Suppose that the current accepted prefix is *M_d_*. The draft model generates a sequence of proposed tokens _, …,_ by sampling from its own next-token distribution with temperature and top-k or top-p sampling applied as in standard decoding. Once this speculative span has been proposed, the target *M_t_* model is evaluated on the concatenation of the prefix and the draft span, and it yields logits for each of the proposed positions.

At the i-th position in the span, the algorithm computes the probabilities of the proposed token under both models. *M_d_* If and *M_t_* denote the normalized probabilities under the draft and target models respectively, the acceptance probability is ______). The algorithm draws a random number and either accepts the draft token with probability

or rejects it. When a rejection occurs, a replacement token is sampled from a residual distribution formed by subtracting the draft probabilities from the target probabilities wherever the target is larger. The decoding then stops using further draft tokens in that speculative group and resumes from the newly accepted token. If all *L* draft tokens are accepted, an additional token may be sampled directly from the target model to regain some of the parallelization benefit.

This procedure has the important property that the marginal distribution of the generated sequence is exactly the same as that obtained by directly sampling from the target model with the same temperature and sampling scheme, so speculative decoding does not alter the qualitative behavior of the generative model.

### KV-Aware Implementation

We considered two ways of integrating speculative decoding with key-value (KV) caching. In the simpler variant, which we mainly used for development, each speculative step runs the draft model on the entire current prefix from scratch, and the target model is run on the prefix plus the draft span. This avoids dealing with partial caching but recomputes hidden states for the full context at every step.

Our main experiments used a KV-aware implementation. In this setting, both draft and target models maintain KV caches that are updated as new tokens are accepted. The initial caches are created by running each model once on the initial prompt. During speculative decoding, the draft model advances its cache incrementally as it proposes each token in the span. The target model is then called only on the draft tokens, making use of its cache for the prefix. After the acceptance phase, the caches must be made consistent with the final accepted sequence: if all tokens are accepted, both caches can simply be advanced; if a rejection occurs, the caches corresponding to accepted tokens are kept, and the residual token or tokens are passed through each model to update the caches appropriately. In some cases, partial KV tensors must be truncated to match the new prefix length.

### Perplexity and Log-Likelihood as Distributional Diagnostics

To verify empirically that speculative decoding preserves the distribution implied by the target model, we analyze the log-likelihood and perplexity of generated sequences under the target. For a completed sequence produced by any decoding scheme, we re-encode it with the appropriate tokenizer and run the target model *M_t_* over the entire sequence. At each position *t* we compute the log-probability of the observed token under the model’s predicted distribution conditioned on the prefix. Averaging these values over the sequence yields a mean log-probability, and its negative gives an empirical negative log-likelihood (NLL). Perplexity is defined as the exponential of this NLL.

Because our sequences are generated with sampling, and because vocabularies such as 6-mer DNA tokens are large, the absolute perplexity values can be high and are best interpreted as cross-entropy estimates rather than intrinsic entropies of the model. For this reason, we focus on differences rather than absolute levels. If speculative decoding faithfully reproduces the target distribution, then sequences generated with and without speculative decoding should have very similar mean log-probabilities and perplexities when evaluated under the target model. We apply this offline scoring procedure to all three models to quantify how closely speculative decoding tracks the baseline target-only decoder.

## Experimental Setup

### Configurations

Our primary metric is throughput, measured as the number of tokens generated per second. For each configuration of model, draft type, speculation window length, temperature, and prompt length, we generate sequences with a fixed number of new tokens and record the wall-clock time for the decoding portion of the inference. Model loading and tokenizer initialization are excluded from the timing. Every configuration is run five times with different random seeds, and the results are summarized by the mean tokens per second and the corresponding standard deviation.

The comparative quantity of interest is the speedup of speculative decoding relative to a baseline that uses only the target model with KV caching. We define speedup as the ratio between speculative and baseline throughput. In addition, we track the mean acceptance rate, defined as the fraction of draft tokens that are accepted by the probabilistic rule, averaged across runs.

The speculation window length, *L,* is varied over small integers, typically values between 2 and 7, For a model with a token vocabulary of size |*V*|, this corresponds to possible candidate sequences within a single speculation window. For our GFMs and PFMs, which operate over vocabularies of size and respectively, this results in between and potential sequences, illustrating the rapid combinatorial growth that motivates restricting *L* to small values, which allows us to observe how more aggressive speculation impacts acceptance and throughput. The vocabulary sizes for DNAGPT, ProtGPT2 and Progen2 are ∼ 16,000, 50,257 and 50,400 accordingly. Temperatures are varied over a modest range, usually from 0.8 to 1.2 in steps of 0.1. Lower temperatures concentrate probability mass on high-likelihood tokens and tend to increase the agreement between draft and target, whereas higher temperatures encourage more exploratory sampling and can reduce agreement. For DNAGPT we additionally vary the prompt length in tokens to test whether longer contexts affect relative speedup; for ProGen2 and ProtGPT2 we retain representative prompts and focus on the interaction between *L*, temperature and draft depth.

For truncated drafts in ProGen2 and ProtGPT2, we vary the number of retained layers to explore the trade-off between draft cost and similarity to the target. In ProtGPT2, where no separate small model is available, all drafts are truncated, and the effective depth is an important free parameter. All experiments are carried out on a single NVIDIA A100 40GB GPU with mixed-precision settings that enable TensorFloat-32 matrix multiplications and optimized convolution kernels. Absolute throughput values depend on hardware and implementation details, but our comparisons are always within the same device and environment.

In parallel with these efficiency experiments, we perform perplexity analysis as described in Methods. For each configuration we sample multiple sequences using both the baseline and speculative decoders and then rescore all generated suffixes with the corresponding target model to compute mean log-probability, NLL and perplexity. This allows us to confirm that any speedup we observe does not come at the expense of drastic changes in the target-model view of sequence plausibility.

### Prompts and conditioning tokens

All benchmarks use identical prompts for the baseline and speculative decoders. For DNAGPT, we prepend each genomic window with the human reference token <R> (or <HUMAN_REF> in the public implementation) followed by a DNA prefix drawn from the hg38 reference, which is then tokenized into non-overlapping 6-mers. For ProGen2, we follow the recommended prompting scheme from Nijkamp et al. 2023, including family and function tags and using 1M as the generation start token when appropriate. ProtGPT2 experiments use unconditional amino-acid generation with the standard start-of-sequence token from the released checkpoint. A complete list of text prompts and their tokenizer-level representations for all models and experiments is provided in Supplementary Table S1.

## Results

For DNAGPT we first examine the effect of prompt length on speculative speedup. We observe speedup as a function of prompt length for different speculation window sizes, demonstrating that the relative gains of speculative decoding are largely independent of prompt length (**Figure 1A**). Whether the prompt consists of a short tag sequence or a longer context approaching the maximum allowed by the model, the speedup remains in a similar range for each value of *L*. This suggests that, once KV caches are used, the main benefit of speculative decoding arises from verifying multiple future tokens per target forward pass, rather than from changes in the amount of past context. When we look at speedup as a function of *L* for different temperatures, a clear pattern emerges. Small to moderate values of *L*, such as 2 to 3, consistently yield the best throughput improvements. Increasing *L* further reduces the acceptance rate, because the draft and target models must agree on longer spans of tokens, and at some point the cost of rejected tokens outweighs the savings from parallel verification. Temperature also modulates this effect. Higher temperatures flatten the distribution and increase agreement between the models and therefore produce higher acceptance rates and larger speedups, while lower temperatures create bigger differences in the models’ distributions and make disagreements more common, particularly for larger windows. Across the explored settings, DNAGPT achieves peak speedups of around 1.25 to 1.35 times the baseline (**Figure 1B**).

**Figure 1:**
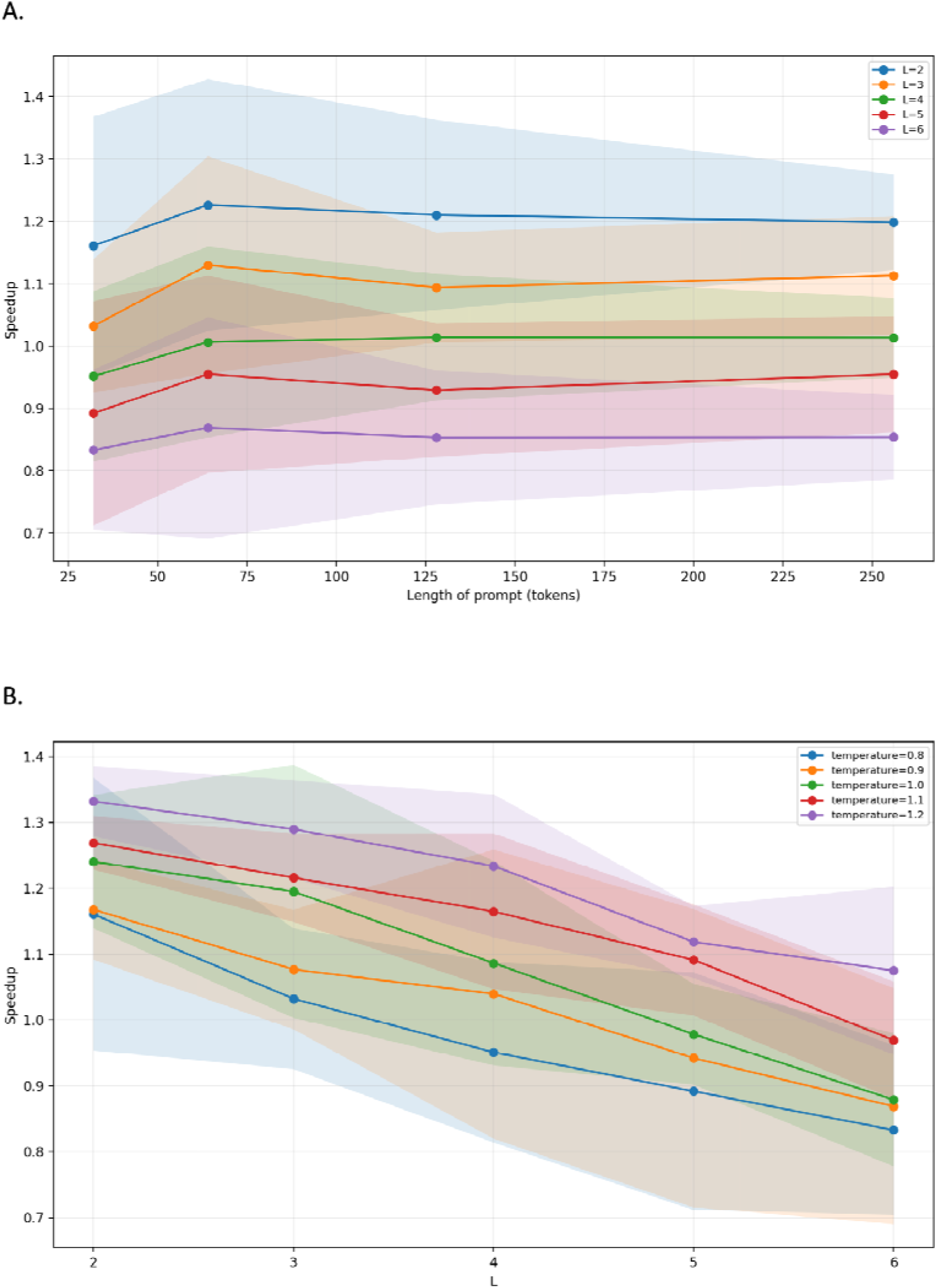
Speculative decoding speedup for DNAGPT. **A**. Mean throughput speedup over the DNAGPT-3B baseline as a function of speculation window size *L*, shown for sampling temperatures *T* = 0.8-1.2; shaded regions denote variability across multiple experiments. **B**. Mean speedup as a function of prompt length (32, 64, 128, 256 tokens) for different speculation window sizes *L*.

In the ProGen2 experiments we compare two kinds of drafts: the separately pretrained progen2-small model and truncated drafts obtained from progen2-xlarge by retaining only a subset of its early Transformer layers. Both approaches lead to consistent improvements in throughput compared to the baseline (**Figures 2A,B**). The pretrained draft typically exhibits higher acceptance rates, which is expected since it is an independent model with its own learned representation of protein sequence structure (**Supplementary Figure 1**). However, it is also more expensive per token than a very shallow truncated draft. Truncated drafts provide a complementary knob. Retaining only a small number of layers substantially reduces the per-token compute cost but also makes the draft less faithful to the target distribution (**Supplementary Figure 2**). In practice we observe that drafts built from a moderate number of layers, for example the first three or four blocks of the base model, strike a good balance: they are cheap enough to generate speculative spans quickly while being accurate enough to maintain reasonably high acceptance rates, especially at moderate temperatures. The resulting speedups again lie in the range of roughly 1.2 to 1.6 times the baseline (**Figure 2B**), depending on the exact configuration.

**Figure 2:**
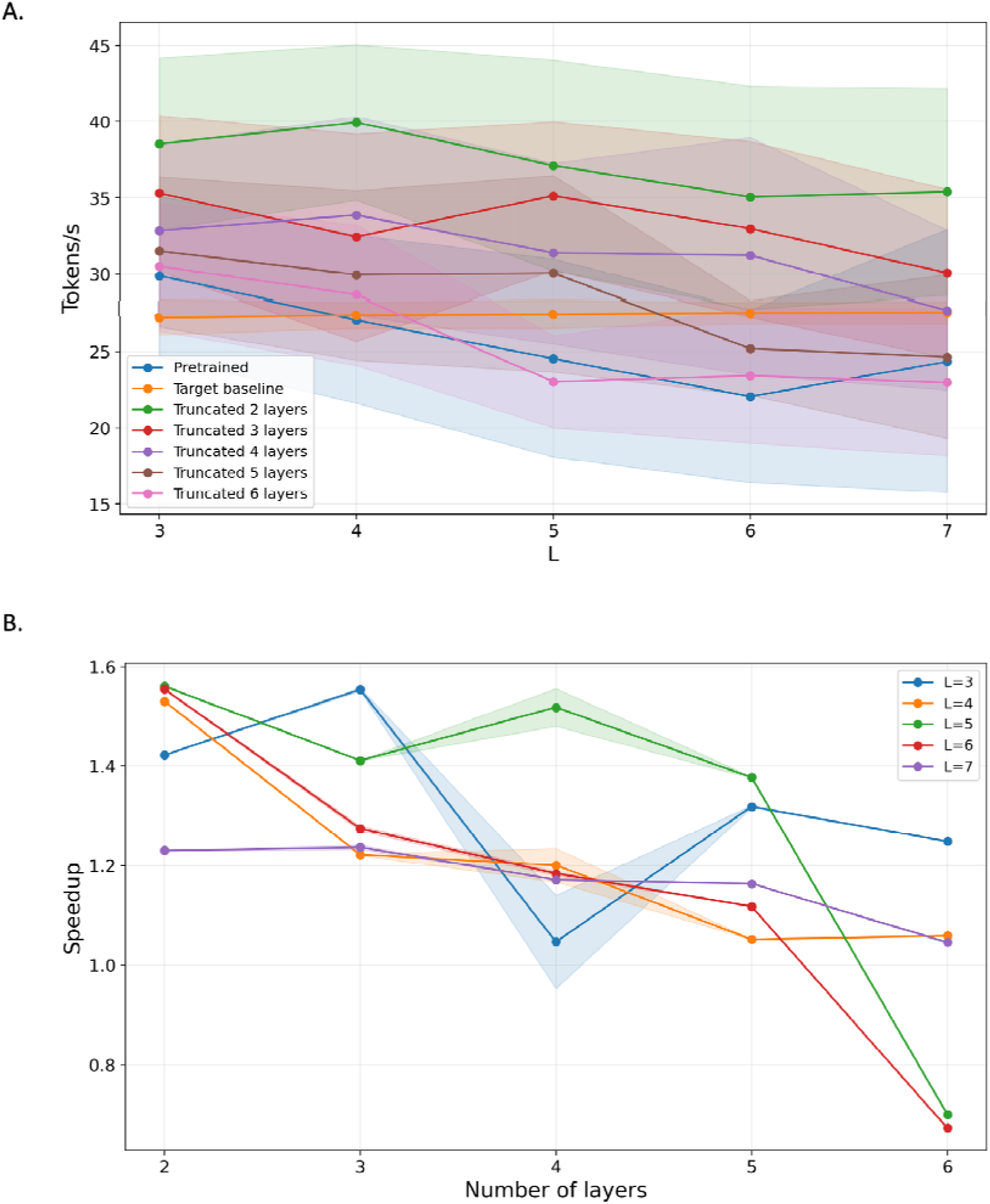
Speculative decoding metrics and speedup for ProGen2. **A.** Mean decoding throughput (tokens per second) for the KV-cached ProGen2 target, the pretrained ProGen2-small draft, and truncated drafts derived from the ProGen2 target with 2–6 early Transformer layers retained, shown as a function of speculation window size L; shaded regions denote variability across repeated runs. **B.** Mean throughput speedup of speculative decoding relative to the KV-cached target baseline when using truncated drafts, plotted as a function of the number of retained layers; curves correspond to different speculation window sizes L= (3 to 7), with shaded regions indicating variability across runs.

**Figure 3:**
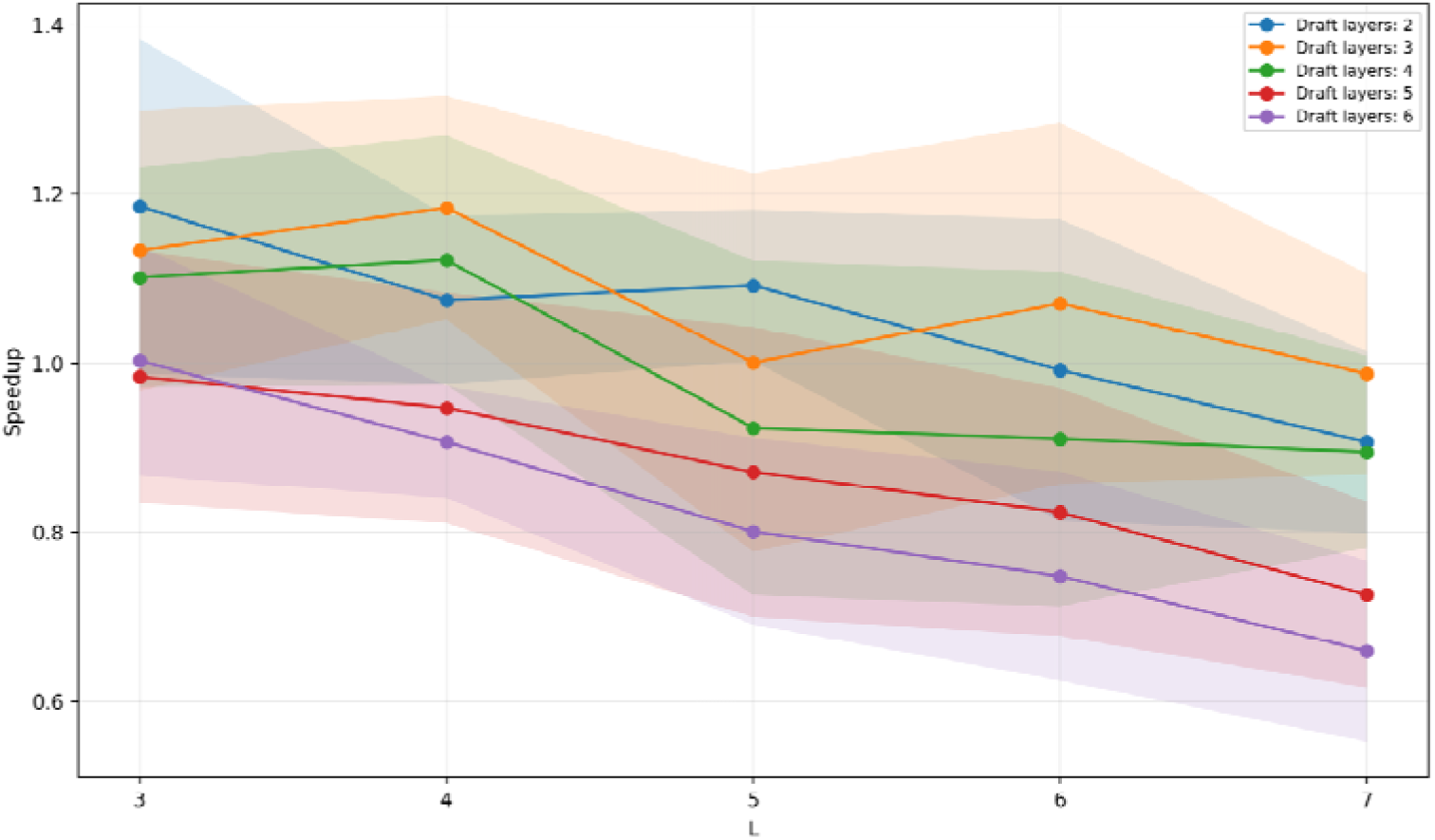
Speculative decoding speedup for ProtGPT2. Mean throughput speedup over the KV-cached ProtGPT2 target baseline as a function of speculation window size L, using truncated versions of the ProtGPT2 target as draft models with 2–6 effective Transformer layers; shaded regions denote variability across repeated runs for multiple configurations.

ProtGPT2 provides an interesting test case because there is no officially released smaller variant that could be used as a draft. Truncated drafts resolve this issue. By constructing drafts that retain only a small number of blocks from the full ProtGPT2 Transformer stack, we significantly reduce the draft’s per-token compute cost while keeping its tokenizer and embedding space identical to those of the target model. When we vary the effective draft depth, we find that very shallow drafts with two to four layers provide clear speedups, whereas deeper truncated drafts with five or six layers have smaller or negligible benefits because their cost approaches that of the full model. For each depth, the optimal speculation window is again small to moderate, mirroring the behavior observed with DNAGPT and ProGen2. Under the best configurations, ProtGPT2 achieves speedups comparable to the other foundation models, typically around 1.15 to 1.2 times the baseline throughput.

### Perplexity Analysis Across Models

Perplexity analysis provides an additional check that speculative decoding truly preserves the behavior of the target model. We extracted perplexity values from the full set of experiments completed and presented in this paper. For DNAGPT we compare the mean log-probability and perplexity of suffixes generated by three decoders: the baseline DNAGPT-3B decoder with KV caching, the speculative pipeline using DNAGPT-0.1B as draft and DNAGPT-3B as target, and the draft-only decoder. When all sequences are rescored under DNAGPT-3B, the baseline and speculative pipelines exhibit nearly identical mean log-probabilities, with differences typically within a few tenths of a nat. Perplexity ratios are close to one across temperatures and speculation windows. The speculative decoder closely tracks the baseline target perplexity, confirming that our implementation preserves the target distribution. We also want to note that sequences generated by the draft-only decoder have slightly lower perplexity under the target model than sequences from the target decoder itself. While this might appear surprising at first glance, it does not contradict the correctness of speculative decoding. Our offline scoring always uses the canonical target distribution (logits at temperature 1, no top-p) to evaluate sequences that were generated under a temperature- and top-p-modified distribution. The small DNAGPT-0.1B draft model is more conservative and tends to sample high-probability k-mers, whereas the DNAGPT-3B target, at the same decoding settings, explores lower-probability tokens in its tail. As a result, the draft-only sequences occupy a slightly higher-probability region of the target distribution, and therefore achieve marginally lower cross-entropy. Conversely, when all sequences are rescored under the draft model, draft-only generations achieve much lower perplexity than sequences produced by the target or speculative decoders (**Supplementary Figure 3**). (**Figure 4**).

**Figure 4:**
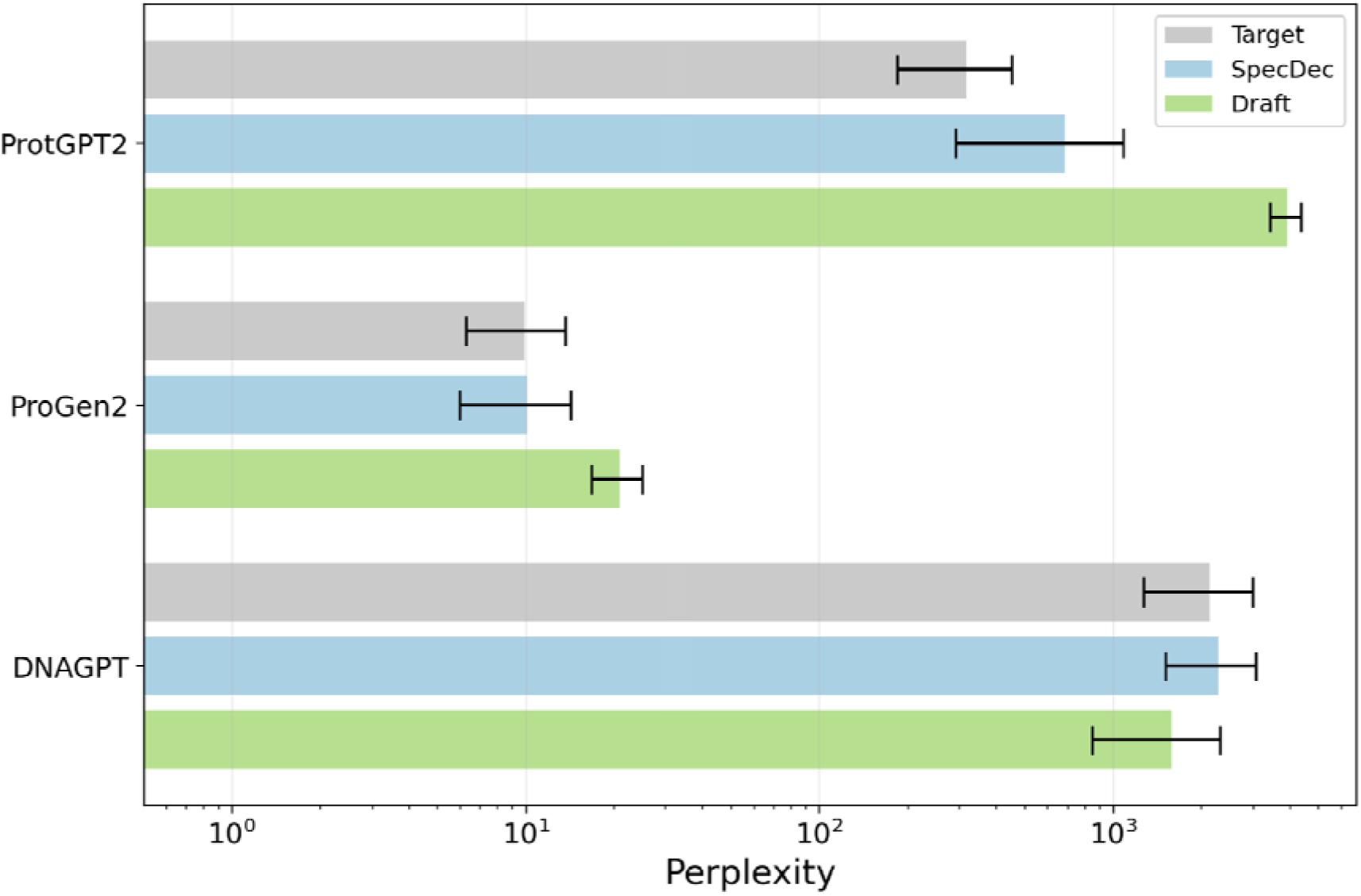
Perplexity comparison across genomic and proteomic models. Horizontal bars show mean suffix perplexity (log scale) when sequences generated by each decoding scheme are rescored under the corresponding target model: ProtGPT2, ProGen2 or DNAGPT. For each model, “Target” denotes target-only decoding, “SpecDec” the probabilistic speculative decoding pipeline, and “Draft” the draft-only decoder; error bars indicate standard deviation across runs.

We observe analogous behavior in the ProGen2 experiments. Sequences generated with ProGen2-xlarge using speculative decoding, whether the draft is ProGen2-small or a truncated version of ProGen2-xlarge, have almost the same target-model perplexity as those generated by the baseline decoder, with small fluctuations that are well within run-to-run variability. Draft-only generations are clearly worse when rescored under ProGen2-base. This confirms that speculative decoding with either a pretrained or truncated draft does not introduce noticeable distributional drift relative to the target model (**Figure 4**).

For ProtGPT2 the situation is similar. Speculative decoding with shallow truncated drafts yields sequences whose perplexity under the full ProtGPT2 model is effectively close to the baseline sequences for matched temperature and sampling settings. Differences are small and do not show any systematic trend as a function of speculation window or draft depth, provided that the draft is not so deep that it becomes nearly as expensive as the target (**Figure 4**).

Taken together, these perplexity results across DNA and protein models support the theoretical claim that probabilistic speculative decoding leaves the target distribution unchanged and demonstrate empirically that our implementation respects this property within numerical precision.

## Discussion

The experiments reported here confirm that speculative decoding is a viable method for accelerating inference in genomic and proteomic foundation models. The relative speedups are more modest than some of the extreme gains reported in certain natural language benchmarks, but they are substantial in the context of biosequence generation, where long sequences and large numbers of runs are common. A 17 to 28 percent reduction in wall-clock decoding time can significantly reduce computational costs in applications such as synthetic promoter design, protein variant exploration or large-scale in silico mutagenesis.

We show that truncated draft models are very suitable components of this process. By reusing embeddings, early layers and the output head from the target model, they provide a form of structural distillation without any additional training. They are easy to construct from any GPT-style architecture that exposes its stack of blocks, and they maintain tokenizer compatibility by construction. Future work could refine this idea by fine-tuning truncated drafts with explicit knowledge-distillation objectives so that their logits more closely approximate those of the full model, which should improve acceptance rates and therefore speedup.

It is also worth exploring the use of state-of-the-art genomic language models such as Evo2 (Brixi et al. 2025) within a speculative decoding pipeline, including low-level integration with their Hyena-style CUDA kernels to correctly exploit block forward passes and state manipulation in long-context sequence modeling for DNA.

Our study emphasizes throughput, leaving the evaluation of downstream biological metrics such as structural plausibility, functional prediction scores, and experimental validation, for future work. The theoretical guarantee that speculative decoding preserves the target distribution in addition to the perplexity analysis results, strongly suggest that downstream behavior will be unchanged, but it is still valuable to verify this empirically on concrete tasks. In addition, our experiments concentrate on unconditional or lightly conditioned generation settings, and additional work is needed to characterize speculative decoding in more complex tasks such as conditional sequence design or sequence-to-sequence translation between biological modalities.

## Code and Data Availability

All relevant code and results can be found on GitHub at this link: https://github.com/Georgakopoulos-Soares-lab/BioSpecDec

## Acknowledgements

Research reported in this publication was supported by the National Institute of General Medical Sciences of the National Institutes of Health under award number R35GM155468 and start-up funds awarded to I.G.S.

## Supplementary Figures

**Supplementary Figure 1:**
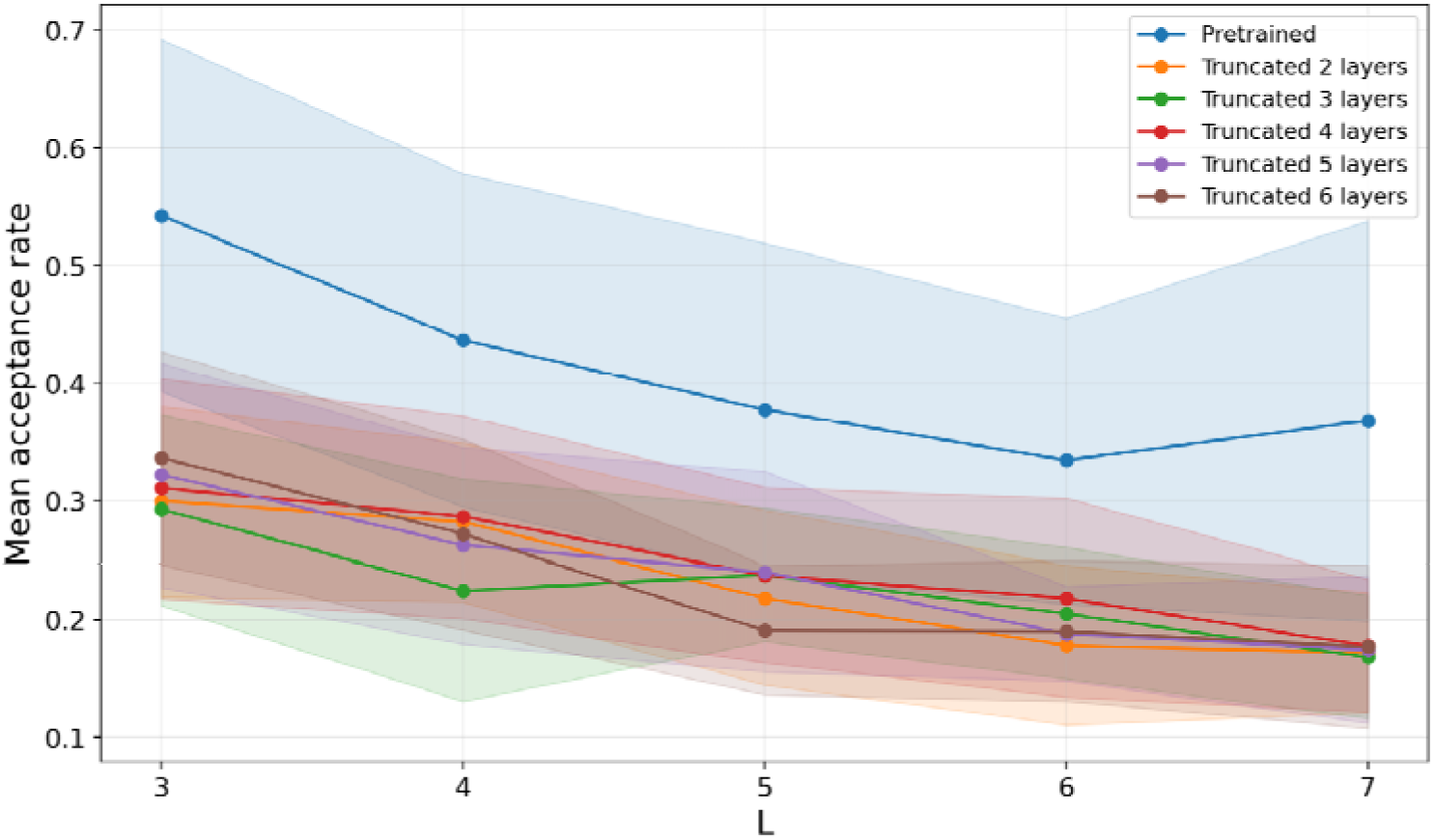
Mean acceptance rate for ProGen2 speculative decoding across draft architectures. Mean draft-token acceptance rate as a function of speculation window size *L* for ProGen2, comparing a pretrained ProGen2-small draft with truncated drafts derived from the ProGen2-xlarge target using the first 2–6 Transformer layers. Lines show the mean over runs and shaded regions indicate variability across configurations.

**Supplementary Figure 2:**
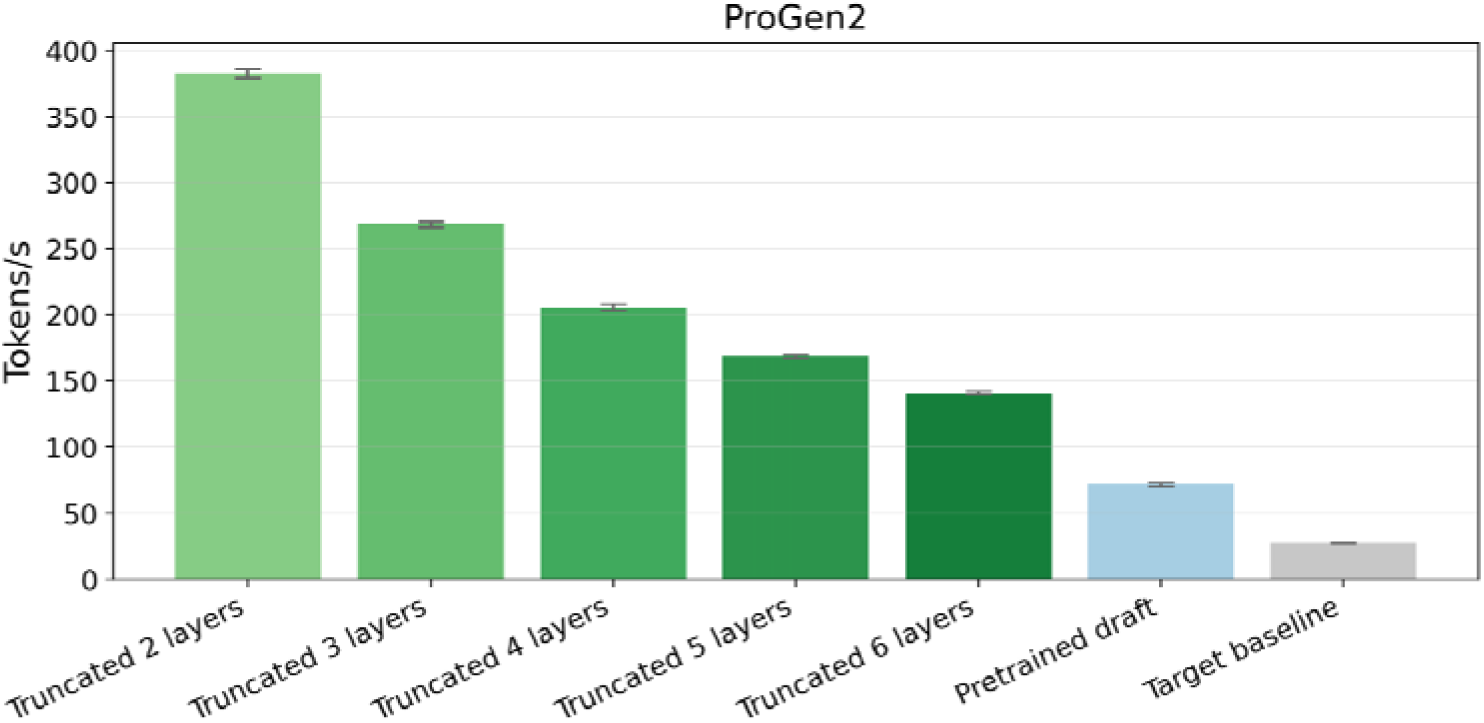
Throughput of ProGen2 speculative decoding for different models. Mean decoding throughput (tokens per second) for ProGen2 when using truncated drafts with 2–6 Transformer layers, a pretrained ProGen2-small draft, and the KV-cached ProGen2-xlarge target-only baseline. Bars show mean tokens/s over repeated runs and error bars denote variability across runs.

**Supplementary Figure 3:**
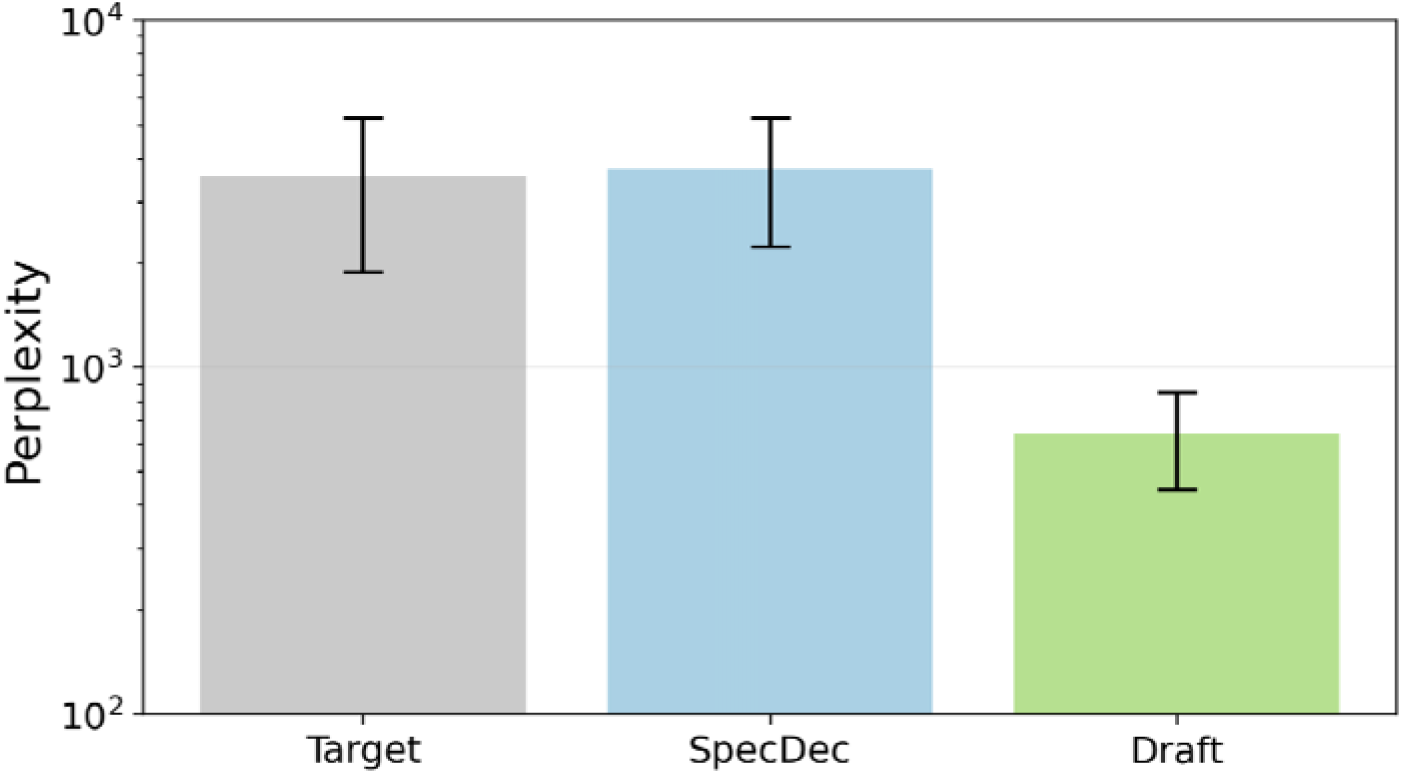
Draft-model perplexity comparison for DNAGPT. Vertical bars (log scale) show mean suffix perplexity when sequences generated by the three DNAGPT decoders are rescored under the DNAGPT-0.1B draft model: “Target” denotes target-only decoding, “SpecDec” the probabilistic speculative decoding pipeline, and “Draft” the draft-only decoder. Error bars indicate standard deviation across generated suffixes.

## Supplementary Tables

**Supplementary Table 1: Sequences / Prompts used for the experiments on the three models.**

**Sequences / Prompts**

